# A general model for genomic traits evolution

**DOI:** 10.1101/2025.10.22.684021

**Authors:** José Ignacio Arroyo, Alejandro Maass, Pablo A. Marquet, Geoffrey West, Christopher P. Kempes

## Abstract

A large body of data indicates that genomic elements, including gene families and entire genomes, exhibit diverse types of evolutionary dynamics. Several models have been developed to describe the dynamics of, for example, genome size; however, a simple deterministic model with interpretable parameters that could be useful for fitting a variety of data remains an open problem. Here, we show that (a continuous form of) the Breeder’s equation for the evolution of quantitative traits/characters leads to a general model that explains the dynamics of genomic elements generated by indel mutations (duplications, insertions, and deletions) and selection. Our framework consists of a general exponential-linear model that predicts at least six different types of dynamical behaviors, including exponential growth and decay. The equations fit data, such as evolution experiments across generations, field observational data of genome sizes across time, and data from phylogenetic reconstructions of ancestral states of genome size across millions of years of a variety of taxa, including viruses, bacteria, fungi, unicellular eukaryotes, plants, and animals, at both intra- and interspecific levels. To test the universality of the dynamics, we derived a dimensionless equation that enables the (re)scaling and subsequent collapse of all data for growth or decay onto a single universal curve. Thus, our model provides a general basis not only for explaining the dynamics in the size of gene families or whole genomes of populations across taxonomic and environmental scales, but also other traits, providing a foundation for further theoretical developments.

## Introduction

Understanding how genomes evolve is fundamental to explaining the origins of genome diversity and complexity, and ultimately the diversity of life [1]. Genome evolution is a twostep process that involves the generation of changes through mutation and their subsequent fixation in populations, which can occur through random drift or natural selection. The mutations that change genome size are called indel mutations. The mechanisms that generate indels include the duplication, insertion, or deletion of genomic parts, a process that occurs during DNA replication or recombination, for example, through unequal crossing over [2]. Despite mutations often being referred to as “random error”, their occurrence is not uniformly distributed across the genome [3, 4], and their average probability across the genome depends on factors such as temperature [5] or genome size itself [6]. The process of duplication, insertion, or deletion creates a set of related genes called gene families, which are groups of genes that share a common ancestor. Classic examples of gene families include CRISPR-Cas, the immune gene family, and transcription-related genes, which can often be a constraint requiring multiple copies of genes [7–9]. In contrast, genome streamlining through deletions and selection is known to occur in highly stable environments, such as in facultative or obligate endosymbiont bacteria, often leading to slow growth [10]. This process can lead to large gene families with thousands of copies when adaptive selection occurs, for example, in the cases of olfactory receptors or immune genes. Genomic material is not only generated endogenously but also introduced by horizontal gene transfer (HGT). HGT can occur within and between any taxonomic domain, including virus-to-virus. All of these types of indel mutations, which increase or decrease genome size, can vary from a single nucleotide to the entire genome.

After an indel mutation occurs, its fixation in the population depends on its fitness effects and the effective population size [1, 11]. The effects of fitness can be neutral, meaning that having an additional copy of an identical or slightly modified gene could cause no change in fitness and be randomly maintained in the genome or removed by selection, given that having extra copies represents a cost. Alternatively, the change in the number of copies of a gene can have a selective advantage, as having an extra copy of a gene and hence extra proteins can be advantageous. However, over time, the initially identical copies can diverge, and one of the copies acquires a slightly different function, such as being expressed in different stages of ontogeny or in different tissues, which may also be beneficial. Examples of this include the globin gene family, for example, which, after different rounds of duplication, have diversified to transport oxygen in different developmental stages and tissues [12]. Gene loss can also be adaptive. For example, facultative or obligate microbial endosymbionts lose many genes that the host can supply, and ultimately evolve very small genomes [13].

Data on genome size dynamics in the literature are limited to a few model species; nevertheless, a wide variety of dynamical behaviors can be observed (e.g., [14–16]). One of the best examples of data on genome size dynamics comes from the Long Term Evolution Experiment (LTEE) [17]. The LTEE is an experimental evolution of 12 *E. coli* populations that reached 50,000 generations by 2016 [17]. Some of the experiment’s results include an increase in genome size and a significant relationship between fitness and genome size in one of the populations [14]. Other experiments have also been performed in yeast, showing a trend to evolve toward diploidy [18]. Examples in multicellular organisms are scarce, but they are present in maize [19], for example. Also, recent studies have focused not only on aggregated properties such as total genome size, but also on the size (number of copies or copy-number for simplicity) of gene families [20]. In *E. coli*, for example, experiments have shown changes in the number of genes in gene families related to antibiotic resistance, responding to different concentrations of antibiotic [20]. At the interspecific level, data show that the amount of genome information, measured as genome size, has increased through evolutionary time, with a correlation between genome size and organismal complexity, from bacteria to plants and animals [21–23].

Several models have been proposed to explain changes in genome composition and size through time [6, 24–28]. These models differ in their mathematical approaches and predictions. Stochastic models include, for example, processes of birth-and-death to explain the distribution of gene families sizes [25, 29, 30], coalescence theory [31] to explain the distribution of frequency of the presence of genes in individuals of a population, or Markov processes that predict genome streamlining [6], and deterministic models include that explain, for example, the relationship between indel rate and genome size [32, 33]. However, there is relatively little testing of these models against data on genome size dynamics generated by experimental evolution experiments [17], field observations [34], or ancestral reconstructions [35]. Despite all this work, a deterministic, simple, and general first-principles model that could be used to fit data on the diverse evolutionary dynamics of genomic structures created by gene duplication-deletion processes (e.g., gene families) has not been developed.

Here, we provide a minimal, deterministic model for the temporal dynamics of a group of related genes that evolve through processes of indel mutations, such as those affecting gene families or entire genomes. To do so, we use one of the most fundamental equations from theoretical evolutionary biology for the evolution of quantitative traits, Breeder’s equation, which incorporates the major forces of genome expansion and contraction rates described above, coupled with selection. We can fit our model to genome dynamics data and estimate key evolutionary parameters, enabling us to interpret the mechanisms underlying any trajectory of genome size change. Our work provides a foundation for future theoretical developments. Our approach represents an efficient theory, that is, a theory built on the minimal set of first principles and assumptions that satisfactorily explain the data [36], rather than on detailed models aimed at each individual species. This approach allows us to find a universal curve for genome changes over time and to compare the effective parameters of different species.

## Results

### Model derivation

We are interested in a general and coarse-grained approach to changes in groups of related genes that evolve via indel mutations, such as gene families or entire genomes, measured as the number of genes or genome size over time, drawing on classical results from population genetics. To do so, we will argue that genomic components, such as gene families or genome size, are quantitative traits that can be modeled using the same theory developed initially for macroscopic traits, such as body size. This argument is based on the fact that the number of copies of a gene family or whole genome, which can be measured in the number of genes or base pairs (bp) or Mb, depends on the genes themselves, particularly the genes that encode for proteins and enzymes involved in replication and recombination [37]. Notice that units such as the number of genes or base pairs (bp) are discrete, but these units can be converted to megabases (Mb), which is a continuous unit. In particular, we will use the Breeder’s equation, a fundamental theorem of evolution. The Breeder’s equation was originally empirically formulated (Lush 1937) to quantify the change in the average trait of a population in the context of selective breeding to achieve genetic improvements in animals and plants, after selection in generation *t* + Δ*t* compared to before selection in generation *t*. Later, Breeder’s equation was theoretically derived in different ways ([38–40], and found to be comparable to other equations, such as Robertson-Price’s equation, re-expressed in alternative ways [41], and refined [40], to account, for example, for natural populations. The time elapsed from one generation to the next, meaning the time between a parent having an offspring and that offspring having an offspring. For simplicity, we will call the generation time *g*. The Breeder’s equation [42] can be formulated as a discrete equation,

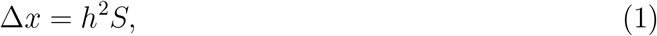

where Δ*x* is the selection response, which is the change in the average trait in offspring 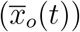 with respect to the entire population 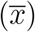, i.e., 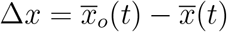. The term *h*^2^ is the narrow sense heritability, which is a parameter that measures the proportion of phenotypic variation in a trait that is attributable to additive genetic variation, and S is the selection differential, which is a measure used in quantitative genetics and breeding to quantify the difference between the average trait value of a selected group of parent individuals 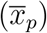 and the average trait value of the entire population 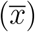, i.e. 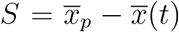. See Table 1 for all terms of this and subsequent equations. Following these definitions, this equation can be rewritten as,

**Table 1.** Relevant variables and parameters of the model.

| Symbol | Definition | Units |
| --- | --- | --- |
| $\Delta x$ | response to selection | bp |
| $S$ | selection differential | bp |
| $\bar{x}$ | average of a genomic trait | bp |
| $t$ | time | generations |
| $h^2$ | heritability | dimensionless |
| $\bar{x}_o$ | average of genomic trait in offspring individuals | bp |
| $\bar{x}_p$ | average genomic trait in parental individuals | bp |
| $g$ | generation time | $\frac{1}{\text{generations}}$ |
| $\sigma_a^2$ | additive genetic variance | bp |
| $\sigma_g^2$ | genetic variance | bp |
| $\sigma_p^2$ | phenotypic variance | bp |
| $\sigma_e^2$ | environmental variance | bp |
| $\sigma_d^2$ | dominance genetic variance | bp |
| $\sigma_i^2$ | interactive genetic variance | bp |
| $\mu$ | mutation rate | $\frac{\text{bp}}{\text{site} \times \text{generation}}$ |
| $k$ | evolutionary rate | $\frac{\text{bp}}{\text{site} \times \text{year}}$ |
| $p$ | probability of fixation | dimensionless |
| $\lambda$ | $2nv_s$ for asexual, $4nv_s$ for sexual populations | number of loci |
| $n$ | number of loci | number of loci |
| $v_s$ | ”spread” parameter in the fitness(-trait) function | |
| $a$ | $\lambda k / \sigma_p^2$ | $\frac{\text{bp}}{\text{generations}}$ |
| $b$ | $\lambda k \bar{x}_p / \sigma_p^2$ | $\frac{\text{bp}}{\text{generations}}$ |
| $N_e$ | effective population size | individuals |
| $s$ | selection coefficient | dimensionless |

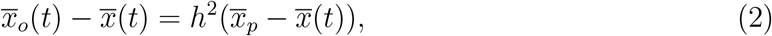

For example, if the average mass of an animal is 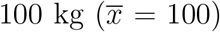, and a fraction with the highest weight is selected for reproduction, which has a weight is 120 kg 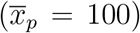, and the heritability *h*^2^ is 0.5 (which as a practical approximation can be measured as the correlation between the weight of offspring and parents), the the predicted response is Δ*x* = 0.5 × (120-100)=10 kg, then the average weight of the offspring will be 110 kg 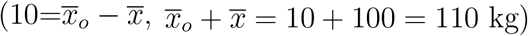.

In equation (2), *h*^2^ is constant and 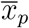 does not depend on time, as is the trait value in generation t before selection. The average trait value of the population of generations *t* and (*t* + *g*), i.e. 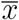, and the average trait value of the population of *t* + *g*, i.e.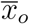 depend on time. Given that 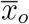 is the average trait in generation *t* + *g*, then 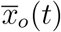 can be rewritten as, 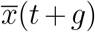, the we can rewrite equation (2) as,

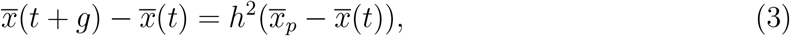

The term *h*^2^, the realized heritability (narrow sense), is the proportion of phenotypic variation due to additive genetic variation and can be expressed as 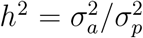, where 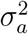 is the additive genetic variance (the total effect on a trait originating from one or more gene loci), and 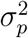 is the total phenotype variance. The additive genetic variance 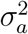 is part of the decomposition of the genetic variance: 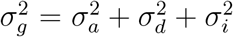, where 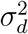 is the variance due to dominance effects, 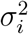 is the variance due to interactive effects. If assuming that 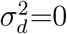 and 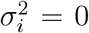, we have 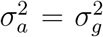 and 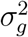, then 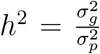. The phenotypic variance can also be decomposed into genetic and environmental 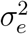): 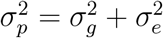 then 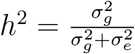.

This last expression can be simplified even further. The genetic variance depends on the mutation rate [43–46]. The mutation rate includes not only indels, generated not only by replication or recombination but also by gains via horizontal gene transfer -HGT-, so this equation would also apply above the species level, since HGT occurs between phylogenetically distant species. The relationships between genetic variance and mutation rate is 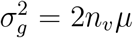 for haploid/asexual [46] and 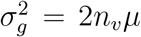 for diploid/sexual [44], where *n* is the number of loci, *µ* is the mutation rate, and *v*_*s*_ is a spread parameter of the exponential quadratic fitness function. As can be seen from the previous definitions, the two expressions differ by a factor of 2 in sexual populations; hence, in both cases, the term is a set of constants times the mutation rate. For simplicity, to define generically a term for genetic variance for both asexual and sexual populations, we will define the term *λ*, then the general expression for genetic variance would be: 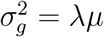, where *λ* = 2*nv*_*s*_ for asexual, *λ* = 4*nv*_*s*_ for sexual populations. It is also known that the practical definition of evolutionary rate (k) is related to the mutation rate (*µ*) and generation time (*g*): 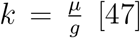, then *gk* = *u*. It is worth mentioning that the expression for evolutionary rate should not be confounded with the similar but different expression for probability of fixation, such as Haldane’s (approximated) equation: *p* = 4*N*_*e*_*µs* (where *N*_*e*_ is the effective population size, *s* is the selection coefficient)[11].

Taking all these together, and using the general expression for genetic variance and mutation rate, 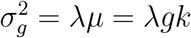, the term heritability can be reexpressed as 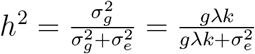. Replacing this expression for *h*^2^ in our previous expression of equation (3), we have,

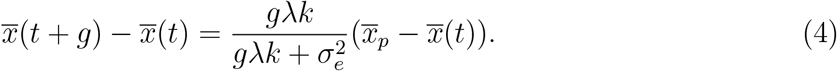

Multiplying both sides of equation (4) by, 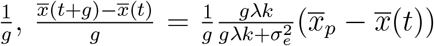 and taking the limit when (g is small - which is the common approach in population genetics theory to go from discrete to continuous time [48, 49], that is *g* → 0, 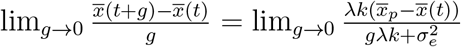,

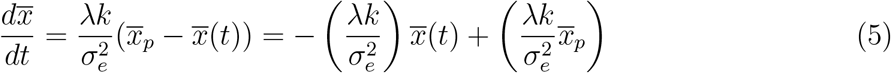

For simplicity, we will rewrite equation (6) as,

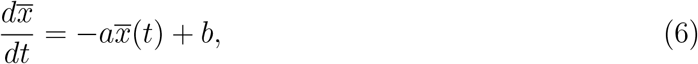

where we define 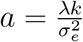, and 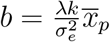. The general solution of this equation is,

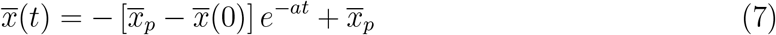

Equation (7) can describe dynamical behaviors of exponential growth or decay. We distinguish at least six different types of behaviors depending on whether the parameter values can be lower, equal, or greater than zero. These different general scenarios are summarized in Table 2 (and specific examples with fitted parameter values will be discussed and shown later in Figure 1). If 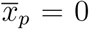, which biologically means that a trait (such as the number of genes in a gene family) is 0–implying *de novo* gene birth–, and i) *a <* 0, there is exponential growth, or ii) if *a* > 0, there is exponential decay with a horizontal asymptote at 0. If 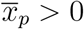, four possible behaviors are possible. If *a <* 0 and iii) 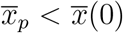 the dynamics starts at x(0) (i.e., the curve intercepts x at x(0)) followed by an exponential growth, and iv) if 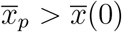 the dynamics starts at x(0) followed by exponential decay intercepting t. If *a* > 0 and v) 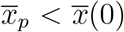 there is an exponential decay with a steady state when t tends to infinity (i.e., a vertical asymptote), and vi) if 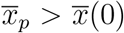 there is an asymptotic growth with a steady state when t tends to infinity (i.e., a vertical asymptote). From making the derivative equal to zero in equation (6), we have that the steady state value of 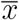, is 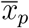.

**Table 2.** Possible dynamical behaviors of the models described by equation (7).

| Parameter values | Dynamic behavior | Example in Figure 1 |
| --- | --- | --- |
| $\bar{x}_p < \bar{x}(0), a < 0, \bar{x}_p > 0$ | Exp. growth intercepting x at $x=\bar{x}(0)$ | A |
| $\bar{x}_p > \bar{x}(0), a < 0, \bar{x}_p > 0$ | Decay intercepting t at $t = \frac{\ln\left(\frac{\bar{x}_p}{\bar{x}_p - \bar{x}(0)}\right)}{a}$ | B |
| $\bar{x}_p = 0, a < 0$ | Exp. growth with intercept at $x=0$ | C |
| $\bar{x}_p < \bar{x}(0), a > 0, \bar{x}_p > 0$ | Exp. decay with horizontal asymptote at $\bar{x}_p$ | D |
| $\bar{x}_p > \bar{x}(0), a > 0, \bar{x}_p > 0$ | Growth with horizontal asymptote at $\bar{x}_p$ | E |
| $\bar{x}_p = 0, a > 0$ | Exp. decay with horizontal asymptote at 0 | F |

**Figure 1:**
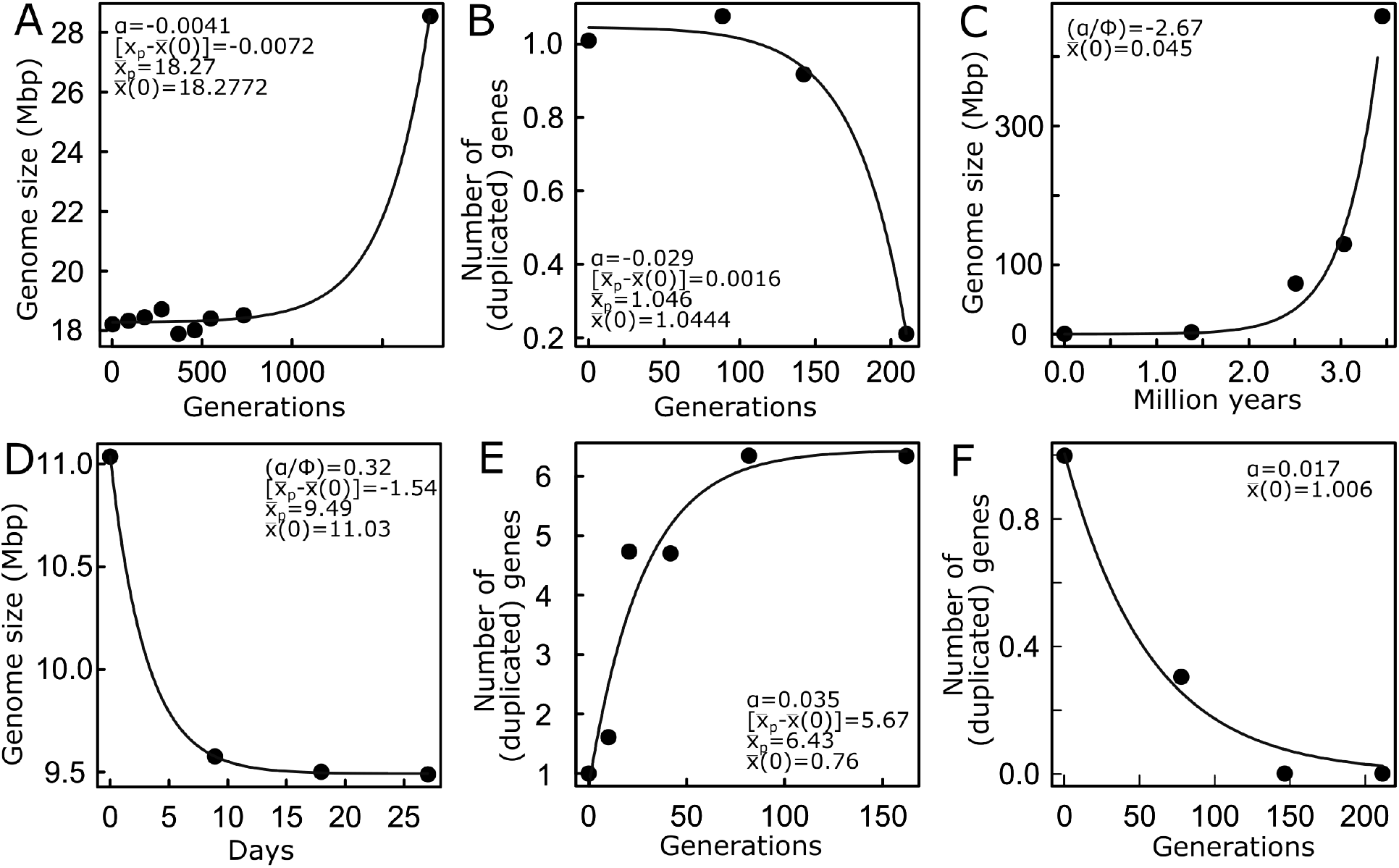
Examples of the dynamical behaviors of the number of genes in a gene family or genome size. A) genome size in yeast (Gerstein et al. 2006), B, F) average number of duplicated genes in the genome in C. elegans (Farslow et al. 2015), C) genome size from bacteria to mammals (Sharow et al. 2006), D) genome size in a plant RNA virus (Willemsen et al. 2019), E) number of antibiotic resistance duplicated genes in E. coli (Pereira et al. 2021).

Notice that in equation (7), time *t* is in generations, but by multiplying by a factor *ϕ* corresponding to the time in other units per generation (e.g., minutes/generations), time can be measured in any other unit, and there would be a new parameter multiplying *t*; 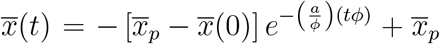, where the term *tϕ* would be time measured in minutes, for example.

#### Dimensionless form

A classic method for exhibiting and testing the generality of an equation is to express it in terms of rescaled dimensionless variables, which predict that a plot of all of the data collapses onto a single “universal” curve [e.g., [50]]. To derive a dimensionless for equation (7), given that at both sides of the equation we have units of, for example, base pairs (bp), we simply divide each side of the equation by 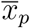, we define dimensionless time as *τ* = *at* for time measured in generations or *τ* = (*a/ϕ*)*tϕ* for time measured in other units such as minutes, after dividing both sides of equation (7) by 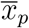 and after rearranging we can rewrite this equation as, 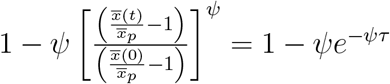,

or simplified as,

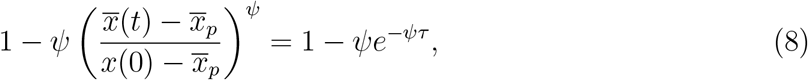

where *ψ* represents a value that can be 1 (if 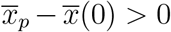, *a* > 0) or 1 (if 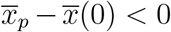, *a <* 0). If *ψ* = 1, the curve describes asymptotic growth, and if *ψ* = −1, the curve describes asymptotic decay.

For the particular case when 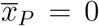, implying 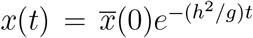, the dimensionless equation would be, 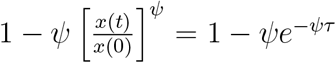, where *ψ* represents a value that can be 1 (if (*h*^2^*/g*) > 0) or −1 (if (*h*^2^*/g*) *<* 0). If *ψ* = 1, the curve describes asymptotic growth, and if *ψ* = −1, the curve describes asymptotic decay. Equation (8) can be derived through different rearrangements besides dividing the original equation (7) by 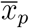, but also by dividing by 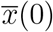 or simply by subtracting 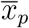 and rearranging.

#### Fitting the model to data

To test the fit of data to equation (7), we collected examples from the literature from experimental and observational studies, for both number of base pairs and genes through evolutionary time, for diverse taxonomic groups, from viruses to mammals (see Methods). Among all the time series in our dataset of dynamics of gene copy number or genome size found in the literature, most of them fitted significantly to equation (7). Some examples are shown in Figure 1, and the remaining fits, together with the estimated parameters, and goodness-of-fit statistics are provided in the Supplementary Material (https://www.dropbox.com/scl/fi/lb83mi3zoq2xig7p39lx4/SM_modgenomeevol_arroyo25.xlsx?rlkey=82fz19wqsuqpvq8psj7isk1l8&st=h6lp80do&dl=0). Each of the specific examples shown in Figure 1 corresponds to a case of all the possible regimes described in Table 2, which include cases of growth and decay (Figure 1). Among the examples, interesting cases include the LTEE experiment in *E*.*coli* [14], experimental evolution in yeast[18] where there is a tendency to evolve toward or remain diploid from either haploid, diploid, or tetraploid populations (Figure 1A), and cases of increase in copy number in *E. coli*, for example, in antibiotic resistance genes in response to different concentrations of antibiotics. There were cases that did not fit any of the general equations, meaning that the change in gene family or genome size was invariant, indicating neutral evolutionary dynamics. We do not show examples of this type here, as we focused on evolutionary dynamics driven by both (indel) mutation and selection. We scaled the data of curves representing genome size dynamics for examples in Figure 1 and others in our database, using both increasing and decreasing patterns, employing the dimensionless equation (8) for data collapse. This equation predicts that decreasing curves should converge to a single curve with an intercept of 2 on the y-axis, and decrease exponentially and asymptotically toward 1 (Figure 2, upper panel). On the other hand, increasing curves should collapse into a single curve with an intercept at zero and grow asymptotically toward 1 (Figure 2, lower panel). In both cases, curves with high goodness-of-fit collapsed into single universal curves (Figure 2), demonstrating the universality of the model, despite the idiosyncrasies of the particular species or evolutionary paths that determined different parameter values.

**Figure 2:**
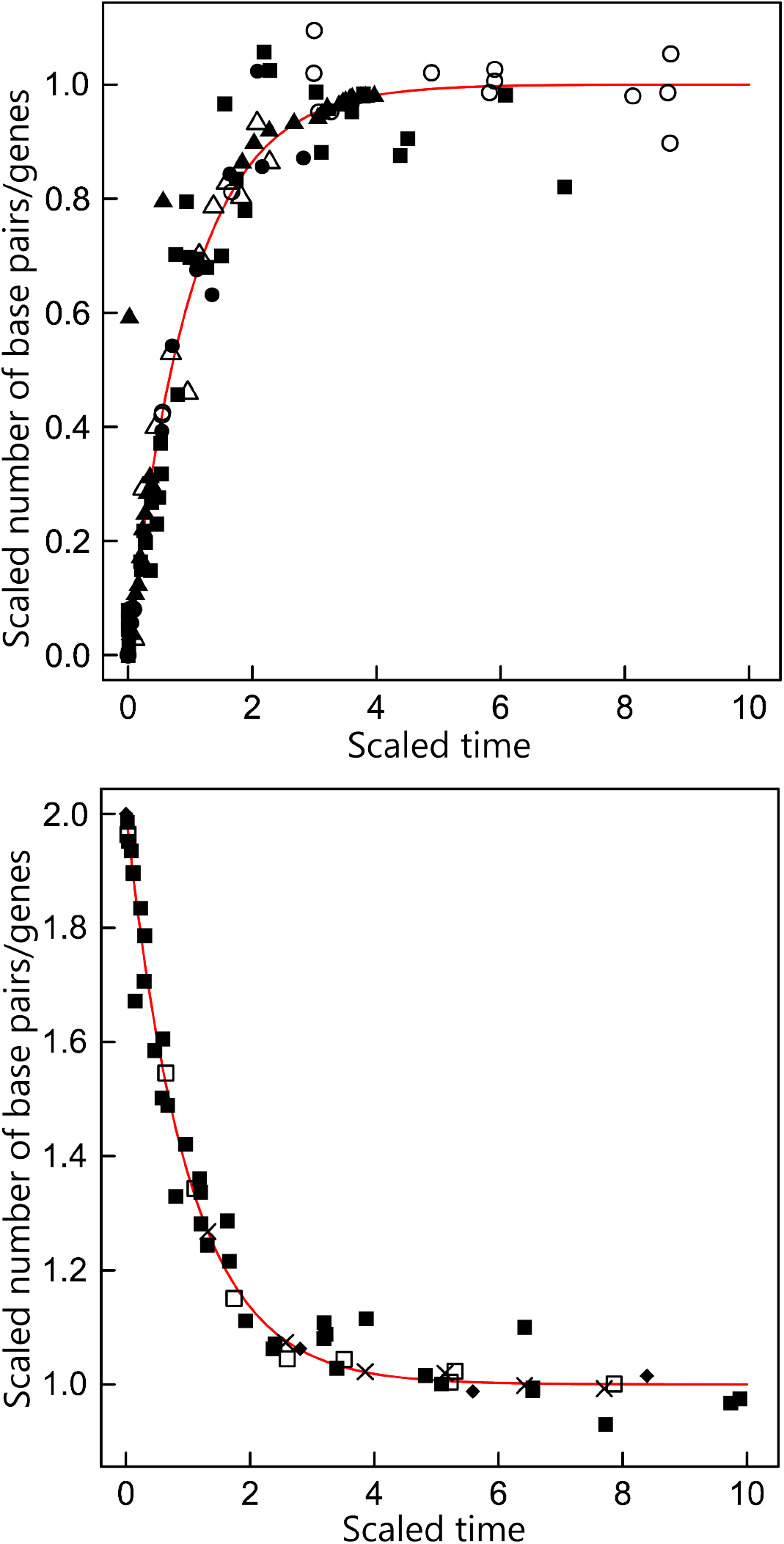
Examples of data collapse. Empty symbols correspond to the number of genes, and filled symbols to the number of base pairs, ×: viruses, □: prokaryotes, ◦:unicellular eukaryotes, △:animals, ⋄:plants. We scaled the data of all significant fits with a high goodness-of-fit (an *R* − *sq.* > 0.8) using the estimated parameters and the dimensionless forms of the equations, for all cases of linear and non-linear selection. When applying this method to different fitted data sets, it predicts a data collapse, i.e., all scaled data should follow the same single ‘universal’ curve predicted by the dimensionless equation. This classic methodology used in physics demonstrates that, despite the idiosyncrasies of the different dynamics, which are probably due to intrinsic (characteristics of the species) or extrinsic factors (environmental context), all of them are governed by the same general equation.

## Discussion

Here we discuss the following: i) previous approaches to model genome size evolution, ii) why we used Breeder’s equation instead of other alternatives, iii) alternatives to Breeder’s equation, iv) limitations of our model, and v) extensions of the model.

Most previous models have focused on formulating a model rather than deriving one, using stochastic models, and often on a single type of mechanism (horizontal or vertical) [6, 24–27]. The approaches and predictions of these models vary widely, including birth-and-death, Markov chains, random walks, agent-based models, and network growth [6, 27, 30, 51, 52]. The predictions encompass the distribution of genes within a gene family, the scaling of rates and size, as well as the stochastic dynamics of genome decay, among others. The missing piece in this variety of approaches was a deterministic model derived from first principles to describe the dynamics of genome structure, including not only genome size but also the size of gene families or any other group of genes emerging from processes of duplication/insertion-deletion.

Although we could have simply derived a phenomenological model for a deterministic dynamics of genomes, our goal was to derive a deterministic model from first principles. For example, using a model analogous to Newton’s heating or cooling law, assuming that genome size (x) change depends on indel mutation rate (m) and size itself but constrained by a minimum or maximum viable size in a specific environment (*x*_*v*_), (given that there is a minimum set of genes necessary for fundamental biological functions, as synthetic cells have demonstrated it [53], and on the other hand there are energetic costs of having too much genomic material.) Following this logic, we could formulate the phenomenological model: *dx/dt* = *m*(*x* −*x*_*v*_).

However, we opted for a first-principles approach here. The importance of developing a first-principles-based theory explaining the origin and evolution of genomic complexity is fundamental, for example, to explain not just the origin of the complexity of genome structures but also transcriptomes and proteomes, as a basis for a theory explain the emerging knowledge of the structure and dynamics of genomes, and especially considering the vast amount of genomics data and upcoming big genome sequencing projects, such as the Earth Biogenome project [54]), and to better understand cancer, as the number of mutations increases with age (i.e., time) [55]. To derive a model from first principles, we used the simplest first-principles approach, Breeder’s equation. We chose to use Breeder’s equation for at least three reasons: i) because it is the most fundamental equation for describing the evolution of a trait. Other equations, such as Lande’s equation, are derived from this equation. ii) Using other equations, such as Lande’s equation or the adaptive dynamics equation, requires more assumptions, such as assuming quadratic polynomials for the fitness functions, some of which do not make sense. For example, assuming stabilizing or disruptive selection (which implies assuming an exponential quadratic polynomial) where only the variance changes and not the mean, results in an equation similar to the one obtained in equation (7), but with this equation. iii) On the other hand, assuming, for example, directional selection, which implies using a linear polynomial, results in a linear dynamics of two parameters, which is statistically indistinguishable from an exponential dynamics, also of two parameters, so there is no way to distinguish statistically between directional selection or another type. Also important to mention is that we used the original form of Breeder’s equation here, but it can be formulated in alternative ways [56], all of which are reducible to a similar mathematical form.

Beyond Breeder’s equation, at least three other slightly more complicated approaches, in the sense that require more assumptions, might have converged to a similar model. These alternative approaches are Lande’s equation and Adaptive Dynamics. Lande’s equation, on one hand, describes the change of the average of a quantitative trait as a function of heritability (which in turn is a function of mutation rate) and the selection gradient, which is derivative of the logarithm of the mean fitness with respect to the trait [57], and is equivalent to Breeder’s equation [58]. The (canonical) equation of Adaptive Dynamics, on the other hand, describes the change of average trait as a function of mutation rate, variance of the trait (of a mutant), (steady-state) population size, and the selection gradient, which is the derivative of the fitness with respect to the trait. As can be seen from the description, both theoretical approaches, evolutionary quantitative genetics and adaptive dynamics, share the commonality that the evolution of a trait depends on the mutation rate and the fitness gradient (i.e., selection) [59]. From these basic expressions, the ultimate form of the equation depends on the selection of the fitness function. Three basic approaches have been described in the literature: directional selection, when one extreme of the traits is favored and can be approached by a linear function. Alternatively, a quadratic approach can be used under stabilizing selection, when an intermediate trait value is favored, or diversifying (also called disruptive) when both extremes are favored [60]. Here, a normal (Gaussian) fitness function ([61], see also [58]). In adaptive dynamics, to conveniently derive a linear differential equation, a quadratic approximation[60] can be used for the fitness function. These two approaches, however, are all phenomenological and have not been derived from first principles yet. The alternative, based on first principles as a function of fitness, is possible for some traits. For example, in metabolic theory, the relationship between fitness and cell size in unicellular organisms, or body size in multicellular ones, has been derived. This derivation predicts a scaling relationship of the form *r* = *r*_0_*m*^*a*^, where *r* is fitness, commonly measured as the per capita population growth rate, *r*_0_ is a constant, *m* is size, and *a* is an exponent that varies depending on the taxonomic group. The limitation of this approach is that although it is possible to establish a theoretical relationship between size and other traits, such as life history traits, and although many traits empirically correlate with size, there is no general model that relates *r* to any trait. Therefore, this first-principles approach is only helpful for a few traits. In addition to the above models, it is possible to use the classical allele frequency models. For example, the model *dx/dt* = *sx*(1 −*x*) − *ux*, whee *x* is frequency of an allele, *s* is the selection coefficient, *u* is mutation rate [62–64]., which is equal in form to Levin’s metapopulation model [65], can be modified to describe the change in the number of alleles or genes.

In our analysis, we found that most cases of both experimental and field data agreed with scenarios of selection, driven by different factors such as antibiotic resistance and stressed medium, among others. In natural populations, the direction of the change —decrease or increase — can be hypothesized to depend on the availability of resources. For example, species that live in hosts, such as symbiont bacteria [66, 67], have reduced their genome because they can utilize specific processed metabolites from the host environment. This process is not exclusive to bacteria, but can also occur in multicellular eukaryotes, for example, in herbivore-plant interactions [68]. There were also many cases of invariant dynamics (see SM) that correspond to instances of just mutation but no selection, where there is a stochastic change in genome size with no trends toward increasing or decreasing. This model has limitations, which are evident from the fact that some patterns are not fitted by the model, such as dynamics with a logistic (S-) shape (e.g., [69–71]), or dynamics with alternative states, such as the transition from haploid to diploid in yeast [72].

This model could be extended in different ways. For example, by integrating this framework with recent developments on the limits of evolutionary rates of traits [73], or empirical studies that show a universal relation between mutation rate and genome size [74] and population size [75].

This model explains previous phenomenological attempts to explain scaling behavior in genomes, such as the scaling of regulatory genes, emerging from coupled exponential dynamics [76], for example. Additionally, this model serves as a basis for further development, as the parameters that define the steady state, for example, are related to other traits such as body size and temperature [77]. Tests of the dynamics of traits for a single lineage are scarce. Probably because the data is also scarce, except for a few traits such as cell size [78] or body size [79].

Not many experimental evolution experiments exist in addition to the LTTE in *E. coli*, as far as we know, there is no other in prokaryotes, but there are a few in unicellular eukaryotes (in yeast and algae)[18, 80]. regarding observational evidence, there are not many time series of long-term observations of a trait, but there must be a few. A well-known example is brain size in humans [81] or at the interspecific level, body size in mammals, for example, [82]. Beyond biology, this model might be applied to any generic process of mutation and selection, such as the growth of companies, which have been argued to be similar to living organisms as they grow and reproduce, and are under selection pressure dictated by markets, consumers, etc. [83]. This strongly suggests that a generic equation or set of equations could be formulated to account for generic processes of endogenous creation/destruction with a feedback structure and exogenous influences.

In conclusion, we formalized the process of evolution of the average size of a genomic component, which originates from processes of duplication/insertion-deletion and selection, commonly including gene families or the entire genome, as well as metabolic pathways, functional categories, and other components. We applied the Breeder’s equation to describe the dynamics of genomic quantitative characters that change due to indel mutations, such as the number of genes in a gene family or the number of genes in a genome, which are traditionally simply measured as genome size. The model is supported by data available in the literature from various gene families and bacterial genomes that have evolved under relatively constant environmental conditions, as well as by data from field observations spanning time. With this study, we demonstrate that a minimal deterministic model can explain the genome evolutionary dynamics of asexual populations. More importantly, constitutes a basis for extending this framework by integrating it with other theories and making new predictions.

## Methods

### Data

We searched for data on the change in the number of genes within a gene family or the number of base pairs (i.e., genome size) over time for a single species or groups of species using Google and Google Scholar. We found articles reporting experimental or observational data, or dynamics inferred from phylogenetic ancestral reconstructions of the evolution of the number of base pairs (genome size) or genes in a gene family in all major taxonomic groups in the tree of life: viruses, bacteria, unicellular eukaryotes, fungi, plants, and animals. Some examples include the Long-Term Evolution Experiment (LTEE) in *E. coli* [14, 84], experimental evolution of yeast genome size under different concentrations of stressors[18], or evolution of the number of duplicated antibiotic resistance genes in response to different antibiotic concentrations [20] (Supplementary Table 1). In many studies, for the same species, there were many populations corresponding to biological replicates that had slightly similar responses. We fitted our model to each population. In our database, we assigned a unique ID to each population, indicating the first author of the article, the year of the publication, the figure and panel, and the specific population, naming it from top to bottom as population 1,2,3, etc., using the following nomenclature “authoryearfigurepanelpopulation”. The total dataset consisted of 31 articles, from which 101 representative time series, each of the size of a gene family or the entire genome, were obtained. Most data on genome size were transformed into Mbp (1*Mbp* = 10^6^*bp*). Some of the genome size data was measured in fluorescence. For simplicity, we just converted the data in the example of yeast, in Figure 1, to Mbp, simply considering that the haploid genome size for yeast is *approx*12 Mbp and for a diploid is *approx*24 Mbp. There were other cases that could also be converted from cell size, for example, as seen in Gallet et al. 2017 [14]. For some cases of the time series in our database of lineages above the species level, it is simpler to transform absolute time to generations because the generation times are not too diverse (e.g., Proboscidea, salamanders), but in other cases, such as a time series from bacteria to mammals, the generation time is too diverse. Also, measuring time for viruses in generations is complicated (but see (Yarwood 1956)[85]), to be transformed to generations, so we transformed time in years or weeks into days.

#### Fitting

To estimate the parameters of the exponential models derived from equation (7); 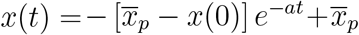, we defined the reduced parameter q, as 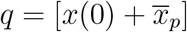. Then equation (7) becomes 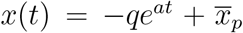. After estimating *q* we can estimate 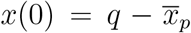. To estimate the parameters *q, a*, and 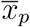, we used the Levenberg-Marquardt (LM) non-linear regression method as implemented in the function “nlsLM” of the “minpack.lm” package [86] in the R language [87]. The starting parameter values that were used are reported in the Supplementary Material. The LM algorithm was run for a maximum of 100 iterations.

To select the best fitting model, we used relative likelihood *L*(*M*_*i*_|*data*), where *M*_*i*_ stands for the likelihood of model i. The relative likelihood was calculated as 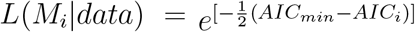, where *AIC*_*min*_ is the AIC of the model with the lowest AIC. The relative likelihood can be interpreted as being proportional to the probability that the *i*th model minimizes the (estimated) information loss [88]. A relative likelihood 2 was considered significant [89].

Data collapse is a classical approach in statistical physics that demonstrates how different curves responding to the same general equation can be plotted in a generic way, with dimensionless parameters all equal to 1, thereby showing that all of them follow the same behavior. Here, for data collapse, we included a few examples of not only fits that were significant but also those with an R-sq. > 0.9.

## Acknowledgements

JIA & CK were supported by SFI and NSF Award Number 2133863. JIA & AM were supported by the Center for Mathematical Modeling (CMM) grant FB210005, which is a Basal fund for centers of excellence from ANID-Chile. AM was supported by the Center for Genome Regulation, which is the Millennium Institute Project ICN2021 044, supported by the Millennium Scientific Initiative of the Ministry of Economy, Development and Tourism (Chile), and Grant Exploración number 13220002.

## Notes

### Competing Interest Statement

The authors have declared no competing interest.

